# Tristetraproline in breast cancer: treat or trick?

**DOI:** 10.1101/2022.05.23.493030

**Authors:** Kropyvko Serhii, Hubiernatorova Anastasiia, Mankovska Oksana, Syvak Liubov, Verovkina Nataliia, Lyalkin Sergii, Lavrynenko Kyrylo, Ivasechko Iryna, Stoika Rostyslav, Rynditch Alla

## Abstract

Breast cancer is one of the most common types of cancer. It is very heterogeneous, hence still complicated to diagnose despite of decades of research. Post-transcriptional regulation of gene expression is crucial for modulation of cell networking and is performed via different regulatory molecules. Tristetraproline (TTP) is RNA-binding protein which binds to AU-rich elements within its target mRNAs and negatively regulates multiple transcripts, including pro-inflammatory and pro-oncogenic. Its expression level correlates with patient outcomes in different types of cancer and is considered as a potential molecular marker. Here we examined TTPs expression level in different molecular subtypes of breast cancer. Our findings show that TTP expression is significantly higher in HER2-enriched breast cancer compared to other types and adjacent tissues. We also investigated changes in the TTPs methylation status under temozolomide and doxorubicin treatment in MCF-7 cell line and found that temozolomide decreased TTPs methylation, which can potentially improve patient prognosis. In contrast, another well-known anticancer agent, doxorubicin, promoted TTPs methylation, which may impair an expected therapeutic effect of this drug.

## 1. Introduction

Breast cancer (BC), despite of numerous studies is still one of the most common types of cancer and remains the leading cause of death from malignant tumors in women. According to International Agency for Research on Cancer, in 2020 the number of BC cases in women reached 24.5% out of overall cancer cases, which corresponds to 2.2 million patients. Moreover, in the last 5 years the percentage of cases was 30.3%, which is more than 7.7 million of patients. In addition, BC also has the highest mortality rate, in 2020 15.5%, of nearly 700.000 women. The BCs well-known feature can explain such high rate of mortality: it is extremely heterogeneous, which complicates the diagnosis and treatment. Traditionally BC is divided in 4 molecular subtypes, based on estrogen, progesterone and HER2/neu receptors expression. In the last decades, numerous studies investigated potential diagnostic and prognostic markers that can increase the specificity of tumor type validation and patient prognosis. Here we investigate tristetraproline (TTP), well-known oncosupressing protein that can be a promising potential marker for BC diagnostics.

Post-transcriptional regulation of gene expression is a crucial step in maintaining of cellular functions and plays an important role in orchestrating proliferation, migration, differentiation, apoptosis and other important events. Despite of cells have multiple regulatory mechanisms, some pathway alterations may lead to dramatic changes, such as malignization. mRNAs possess different regulatory elements, mainly in their 5’ and 3’ untranslated regions that act as binding sites for different regulatory molecules, such as RNA-binding proteins (RBPs). RBPs act as spatiotemporal regulators of gene expression, moderating target mRNAs’ stability and accessibility for translation, as well as their transport to certain compartments for local translation or to processing bodies, thus flexibly regulating cellular network. Whilst changing an expression of single regulatory RBP cell undergoes the alteration of the whole set of its target mRNAs, hence pointing RBPs as worthy targets for cancer research.

Tristetraproline family (TTP, ZFP36 family) is a family of zinc finger RBPs that in humans and other mammals consists of 3 members: TTP itself, encoded by *ZFP36* gene, which is located on 19 chromosome in 19q13.2 region, ZFP36L1 and ZFP36L2, encoded by *ZFP36L1* and *ZFP36L2* genes, respectively. One more member of this family, ZFP36L3, is exclusively expressed in rodent placenta [1]. These proteins contain 3 domains: nuclear export signal (NES) at N-terminus, tandem CCCH zinc finger domain and CNOT binding domain at C-terminus [2–3]. TTP binds to AU-rich elements (AREs) in 3’UTRs of its target transcripts in sequence-specific manner and promote mRNA decay in different ways, such as deadenylation, inhibition of polyadenylation of pre-mRNA or 5’-3’ decay by enhancing decapping [4–7]. Modulation of TTPs functionality can be achieved in different ways: loss/decreasing of expression by *ZFP36* promoter methylation, microRNA-induced mRNA silencing and regulation of protein stability/activity through phosphorylation [8]. TTP protein plays an important role in immunity and cancer, regulating different target mRNAs involved in immune response, cell cycle controlling, proliferation and apoptosis in different manners and is intensively investigated in these fields being a promising onco-suppressor [9–10].

Here we investigate the levels of *ZFP36* expression in patients with different tumor molecular subtypes and its methylation status under the temozolomid and doxorubicin treatment.

## 2. Materials and methods

### 2.1. Total RNA isolation from breast tumor samples

Breast tumor samples were obtained from the National Cancer Institute, frozen in liquid nitrogen immediately after surgery and stored at −80°C. All subjects signed written informed consent. Total RNA was isolated from 0.2-1.5 g tissue by guanidinium isothiocyanate method using the innuSOLV reagent (Analityk Jena) or RNA Go (BioLabTech) in accordance with the manufacturer’s recommendations. Clinical information on the obtained samples is presented in Table 1.

**Table 1.** Clinical information of samples included to the study.

| Histological type of sample | Sample number |
| --- | --- |
| Invasive carcinoma | 53 |
| Adjacent tissues | 13 |
| Normal tissues | 1 |
| Stage |  |
| I | 11 |
| II | 33 |
| III | 7 |
| IV | 2 |
| TNM classification (**t**umor, **n**odus and **m**etastasis) |  |
| T1 | 14 |
| T2 | 36 |
| T3 | 1 |
| T4 | 2 |
| N0 | 37 |
| N1 | 10 |
| N2 | 6 |
| M0 | 52 |
| M1 | 1 |
| Receptor status |  |
| ER<sup>+</sup>PR<sup>+</sup>HER2/neu<sup>-</sup> | 13 |
| ER<sup>-</sup>PR<sup>-</sup>HER2/neu<sup>-</sup> | 12 |
| ER<sup>+</sup>PR<sup>+</sup>HER2/neu<sup>+</sup> | 11 |
| ER<sup>-</sup>PR<sup>-</sup>HER2/neu<sup>+</sup> | 17 |

### 2.2. cDNA synthesis

5-8 μg of total RNA was pre-treated with DNase I (Fermentas, Lithuania), according to the manufacturer’s recommendations, to remove residues of genomic DNA. After that, cDNA synthesis was performed in 20 μl according to the manufacturer’s recommendations. The cDNAs were stored at −20°C.

### 2.3. PCR with fluorescence labeled probes

PCR was performed in 25 μl of mixture containing 0.2 μM of each specific primer and 0.1 μM Taq-Man probe, 1.5 mM MgCl2, 0.2 mM dNTP, 2.5 units DreamTaq DNA polymerase (Fermentas, Lithuania) and the corresponding buffer. Amplification was performed under the following conditions: denaturation - +95°C, 20 sec (in the first cycle - 2 minutes); the time and temperature of the primers reassociation and synthesis were combined: +60°C 1 min, for 50 cycles. Each sample was analyzed as duplicates. The PCR was performed on CFX96 BioRad. The *TBP* gene was selected as a reference based on the analysis of literature sources that showed its appropriateness as a control for the genes expression analysis in breast cancer [11–13]. Primers and probes used for PCR: (For. *TBP* 634-654 5’gtgcccgaaacgccgaatata3’, Taq-Man probe *TBP* 655-676 5’(BHQ1)atcccaagcggtttgctgcggt(FAM)3’, Rev. *TBP* 708-688 5’ccgtggttcgtggctctctta3’); (For. *TTP* 58-79 5’catggatctgactgccatctac3’, Taq-Man probe *TTP* 98-117 5’(FAM)agccctgacgtgcccgtgcc(BHQ1)3’, Rev. *TTP* 177-195 5’ctggagtcggaggggctca3’).

The primers nucleotide positions correspond to the human *TBP* cDNA with the GenBank accession number NM_006277, the human *TTP* – NM_003407.5.

### 2.4. Real time PCR calculations

The following formula was used to calculate PCR results: Exp = 2^(E_ref_^−Ct(ref)^-E_target_^−Ct(target)^), where E_target_ – the PCR efficacy for the target gene, E_ref_ – the PCR efficacy for the reference gene, ^Ct(target)^ – mean cycle value for the target gene, ^Ct(ref)^ – mean cycle value for the reference gene. The PCR efficacy was determined using calibrating curve method. The cycle values for the target and reference genes were determined as the point of intersection of the fluorescence curves with the threshold, above which the fluorescence value is considered significant. The threshold level was set the same for all experiments at100 RFU. Statistical processing of the PCR data was carried out with Origin 2021b software using the Kruskal-Wallis ANOVA method and Mann-Whitney criterion.

### 2.5. Cell culture and treatment

MCF-7 human breast cancer cell line was obtained from RE Kavetsky Institute of Experimental Pathology, Oncology and Radiobiology, National Academy of Sciences of Ukraine (Kyiv, Ukraine). Cells were cultured in Dulbecco’s modified Eagle’s medium (DMEM, Sigma, USA) supplemented with 10% fetal bovine serum (Sigma, USA) in a CO_2_-incubator at 37°C, 5% CO_2_, 95% humidity and reseeded at ratio of 1:5 every 3-4 days.

Cells were seeded on 60 mm cell culture dishes (Nest Biotechnology, China) at concentration of 1 000 000 cells per each. After 24 h of an attachment period, Doxorubicin was added at concentration 0,1, 0,5, 1 μM and Temozolomide at 100, 150, 200 μM. Following 48 h of incubation, cells were trypsynized (Trypsin-EDTA solution, Sigma Aldrich, USA) and counted using the Trypan Blue dye at 0.01% final concentration (Invitrogen, Thermo Fisher Scientific Corp., USA) in hemocytometric counting chamber. The cells were washed 2 times with cold PBS and frozen at −20°.

### 2.6. DNA isolation

DNA was isolated from cells sediments, previously resuspended in lysis buffer (10mM Tris, 0,1 mM EDTA, 0,5% SDS, pH 8.0) and incubated for 15 minutes at room temperature. Then proteinase K was added to the samples (final concentration 100 mg/ml) and incubated for 30 minutes at 50°C. Then, 1 ml of phenol : chloroform : isoamyl solution (25:24:1) was added to each sample, mixed and centrifugated at maximum speed during 15 minutes. The upper DNA-contained phases were replaced to the new eppendorph tubes and the equal volume of chlorophorm was added. Samples were mixed and centrifugated, the upper phases were replaced to the new tubes and 1,5 volume of 99% cold ethanol was added for the DNA precipitation. Then samples were incubated at −20°C for several hours and the DNA sediments were obtained by centrifugation, followed by 2 washing steps. Then the sediments were dissolved in 25 μL of Tris-EDTA buffer.

### 2.7. Bisulphite convertion and quantitative methyl specific PCR (qMSP)

Before bisulfite treatment the concentrations of the samples were measured by the NanoDrop and concentrated samples were dissolve to the appropriate concentration (<250 ng/mcl, to avoid the non-complete DNA convertion). For bisulfite treatment we used EZ DNA Methylation-Lightning Kit (Zymo Research) according to the manufacturer’s instructions. qMSP PCR was performed using HOT FIREPol® EvaGreen® qPCR Mix (Solis BioDyne). The CFX96 Touch Real-time PCR Detection System (Bio-Rad, USA) was used to perform the reaction. Reaction conditions were 95°C for 12 min, 40 cycles of dissociation at 95°C C 20 sec, annealing at 61°C for 15 sec and elongation at 72°C for 10 sec, followed by melting of PCR product from 65°C to 95 °C increment 0,5°C, 5msec, 60 repeats. Methyl Primer Express software and Li LC and Dahiya R. MethPrimer: designing primers for methylation PCRs. Bioinformatics. 2002 Nov;18(11):1427-31. PMID: 12424112 were used for ZPF36 primers MSP primers design. OLIGO ANALYSIS TOOL by Eurofins [https://eurofinsgenomics.eu/en/ecom/tools/oligo-analysis/] was used to analyze primers for formation of self-dimers and cross-dimers in PCR. Primer-BLAST by NCBI was used to find out the possible non-specific targets in *Homo sapiens* genome (without bisulfite convertion). Resulting primers for identification of ZPF36 methylation have the following sequences: Methylated: F 5’ TAGGGTTAGTTAGGTTGCGTC 3’,R 5’ CACCGAAAACCGACTACTTATA 3’, PCR product length 122 bp; Unmethylated: F 5’ TTTAGGGTTAGTTAGGTTGTGTT 3’, R 5’ CACCAAAAACCAACTACTTATAAA 3’, PCR product length: 122 bp.

Alu and Col2A1 genes were used as references [14–15]. The quantification of relative amount of methylated and unmethylated form of genes was performed using 2^Δ Ct^ method, where Δ Ct=Ct (Col2A1) – Ct (gene of interest (methylated or unmethylated). Then the methylation rates were calculated as equation [relative amount of methylated gene] / [relative amount of methylated + relative amount of unmethylated gene]*100%. The changes of methylation relative to the control (untreated) sample were calculated as [% of methylation in control sample/% of methylation in analyzed sample]. The enzymatically methylated human genomic DNA (methylated Control DNA kit, Abcam) was used as positive control in this study.

## 3. Results

### 3.1. TTP mRNA expression analysis in breast cancer samples by quantitative RT-PCR

Previously, Goddio and collaborators analyzed DNA microarray datasets of 295 primary invasive breast carcinomas classified according to the five genomic intrinsic subtypes and on clinical samples, but their clinical samples were divided only in 3 groups: normal tissues, normal adjacent tissues and invasive carcinoma [16]. Taking this to the account, we analyzed relative TTP expression levels in tumor samples of various stages to explore whether it may be a potential marker of breast cancer types. Clinical samples (n=67) were obtained from the National Cancer Institute. Table 1 shows the tumor data of patients whose samples were analyzed in this study. Fig. 1 shows relative TTP expression levels in different BC subtypes and adjacent tissues.

**Fig.1.**
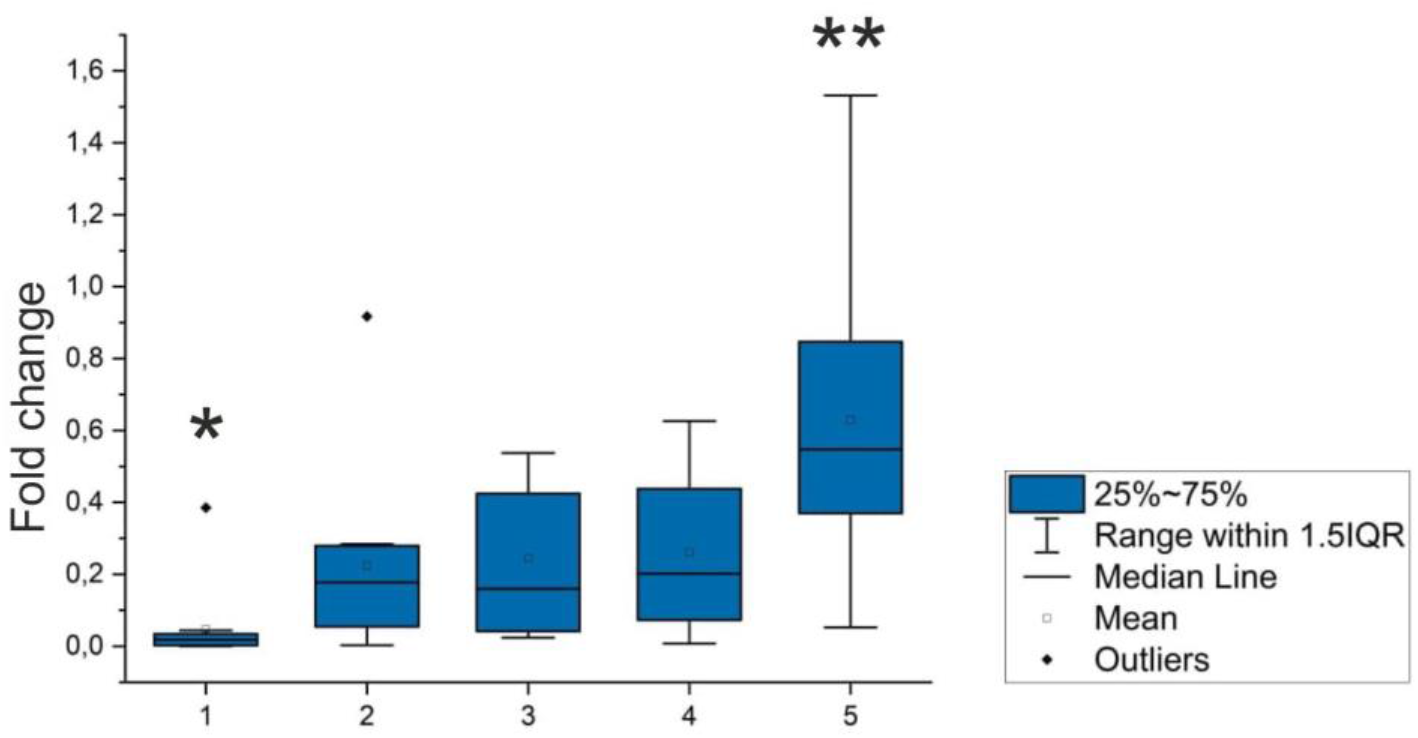
Relative TTP expression levels. 1 – adjacent tissues (n=14), 2 – triple negative type (n=12), 3 – luminal A type (n=13), 4 – luminal B type (n=11), 5 – HER2-enriched type (n=17). * - p<0.05, ** - p<0.01.

The conducted analysis showed that TTP expression was significantly increased in tumor samples compared to adjacent tissues (p<0.05) that nearly comes into agreement with published data. Interestingly, it is also dramatically increased in HER2-enriched tumor samples compared to other tumor subtypes (p<0.05, p-value between adjacent tissues and HER2-enriched p<0.01).

### 3.2. ZFP36 methylation under the Doxorubicin and Temozolamide treatment

In all samples both methylated and unmethylated sequences were amplified. We verified the size of those sequences using electrophoresis in 3% agarose gel and found, that all sizes correspond to the expected (122 bp for methylated and 122 bp for unmethylated) (Fig 2).

**Fig. 2.**
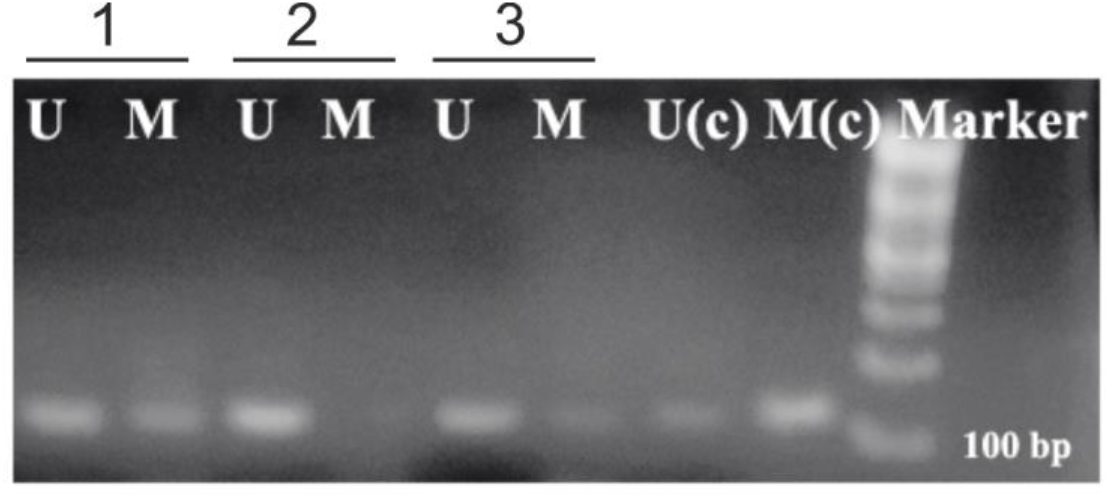
The example of the electrophoresis of qMSP PCR fragments (Temozolomide in different concentrations, 1 - 100, 2 - 150 and 3 −200 μM, (c) - control methylated DNA, marker (Thermo scientific, Gene ruler SM0331)); U – <u>U</u>n-methylated, M – <u>M</u>ethylated

We found that in MCF-7 *ZPF36* is methylated at 11.85%. Studied substances influenced the methylation of this gene in different manner, moreover, this influence depended on the concentration of these chemicals (Fig. 3).

**Fig. 3.**
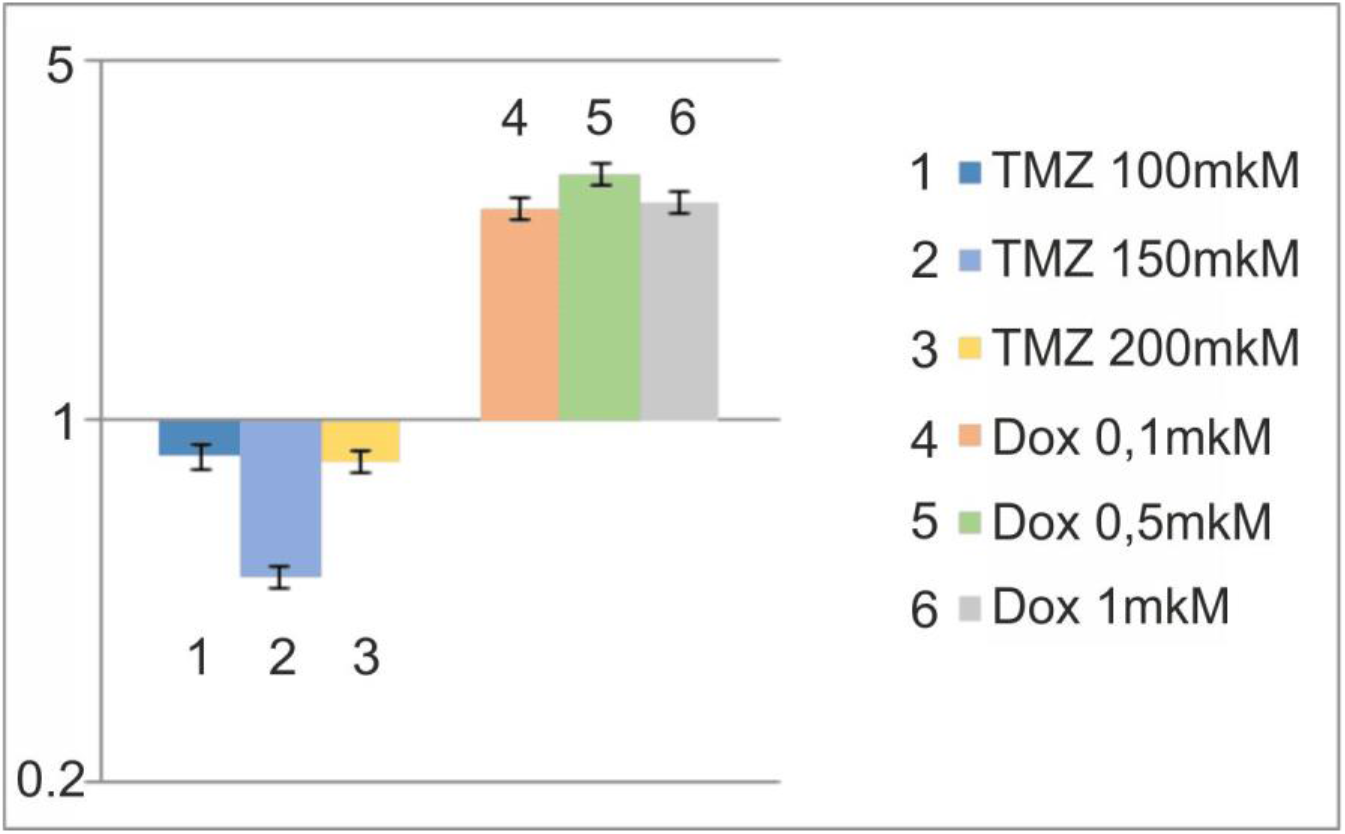
Changes of methylation of ZPF36 after treatment with different concentrations of Temozolomide (TMZ) and Doxorubicn (Dox) in comparison with untreated MCF7 cells (logarythmic scale).

Figure demonstrates the opposite mode of action of the studied drugs on methylation of *ZPF36* in comparison with its methylation in untreated MCF-7 cells. Temozolamide reduces its methylation, and the most effective dose for this effect is the middle concentration, namely, 150 μM.

At the same time, doxorobicin increased the methylation of *ZPF36* gene promoter. The máximum procentage of its methylation, according to our calculations, was 35.4% (after treatment with 0.5 μM of doxorubicin) (Table 2).

**Table 2.** Methylation levels of ZFP36

|  | Control MCF7 | TMZ 100 | TMZ 150 | TMZ 200 | Dox 0,1 | Dox 0,5 | Dox 0,1 |
| --- | --- | --- | --- | --- | --- | --- | --- |
| ZPF36 Methylation | 11,85% | 10,10% | 5,90% | 9,84% | 30,60% | 35,40% | 31,14% |

**Table 3.**
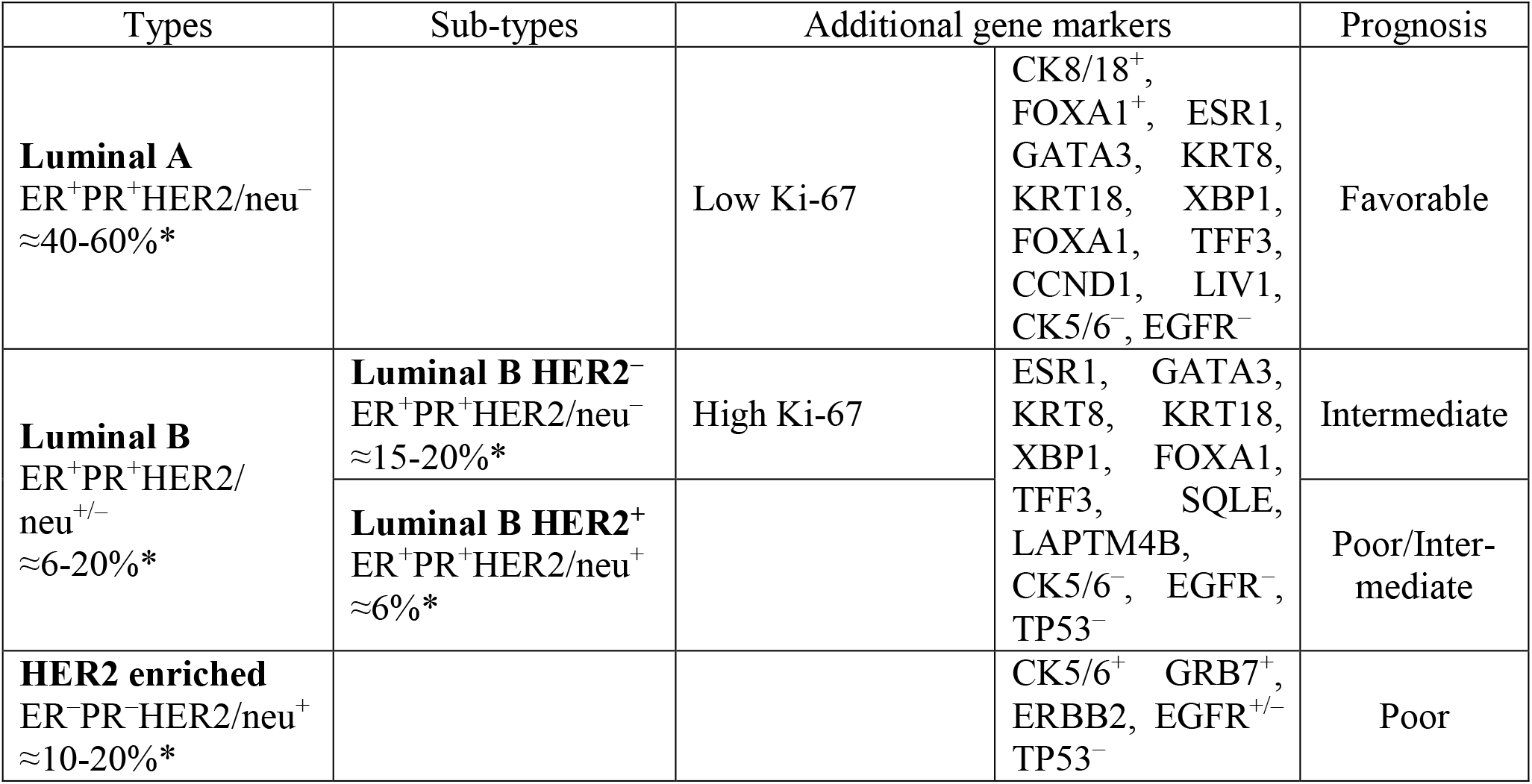
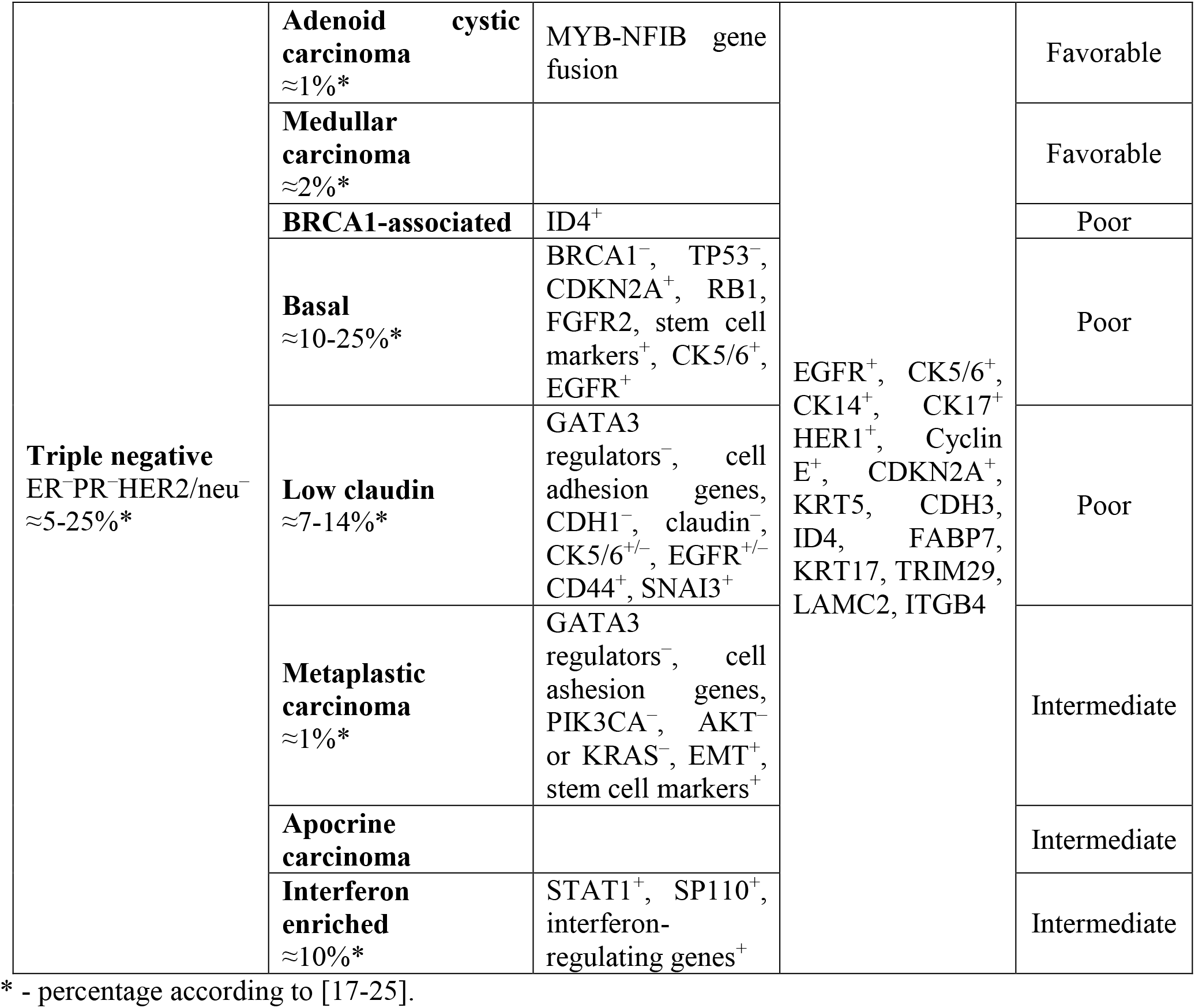
Breast cancer classification: histology, molecular markers and survival prognosis [26]

## 4. Discussion

As mentioned above, breast cancer types are divided into 4 major groups depending on clinical data and specific marker gene expression pattern [17–26]. Molecular types are usually diagnosed using the information about presence or absence of estrogen receptor (ER), progesterone receptor (PR) and HER2/neu receptor expression and further prognosis and therapy strategies are based on this classification. However, decades of research have shown the presence of a large number of subtypes even within these ones, showing that the same type of tumor in different patients can both be more or less malignant and thus have poorer or better prognosis. For example, triple negative BC includes at least 7 subtypes: 3 with poor prognosis, 2 intermediate and 2 favorable ones. Every subtype is characterized by specific marker genes expression, and year to year, their number continues to increase.

Tumorigenesis is characterized by increased expression levels of different oncogenic genes that allow uncontrolled cell division, proliferation and migration, which can be a result of oncogene suppression loss. TTP negatively regulates many oncogenic transcripts and is a promising biomarker since its expression is altered in multiple types of cancer and frequently is associated with clinical features and patient prognosis, moreover, it can modulate the interaction between immune and cancer cells [27–28]. For example, in prostate cancer, patients who developed metastatic disease showed lower TTP expression level compared to those who had localized disease. In patients with radical prostatectomy, prostate cancer with low TTP expression level showed decreased time to biochemical recurrence or metastasis compared to those with high TTP expression level [29–30]. TTP protein and mRNA levels were also decreased in bladder cancer; patients with high expression of its target genes, including those participating in cell cycle control, epithelial to mesenchymal transition and Wnt signaling, had poorer prognosis and more active tumor proliferation compared to bladder cancer patients with lower TTP target genes expression. Interestingly, newly identified by authors doublestranded RNA dsTTP-973 could increase TTP expression level and suppress tumor aggressiveness in vitro and in vivo [31]. Numerous studies investigate TTPs function in breast cancer [8], but not only in the context of ARE-containing mRNAs expression regulation. One interesting study shows that TTP expression level has a significant impact on ERα activity on MCF-7 cells reducing SRC-1 effect on ERα activation. Interaction between TTP and ERα in the same cell line in *vivo* represses ERα transactivation without affecting its mRNA level, suggesting that the repression is implemented exactly via protein-protein interactions [32].

TTP is significantly downregulated in human hepatocellular carcinoma (HCC) and intrahepatic cholangiocarcinoma, which is associated with hepatocyte dedifferentiation and correlates with poor prognosis. Interestingly, in the same study the authors show that TTP also shows a tumorigenic effect and promotes tumor initiation in mice, assuming that it has dual role depending on the tumor formation stage [33]. Surprisingly, our data showed that it significantly overexpressed in HER2-enriched breast cancer samples. Taking to the account that most samples were II grade (33 out of 53 samples) and non-classical reports given above, we suggest that such expression level might be a compensatory mechanism for tumor suppression. Moreover, according to Eliyatkin and coauthors, HER2-enriched subtype accounts only for 15% of invasive breast cancer cases, whilst 50% of cases were diagnosed with luminal A subtype [34]. Our study shows that the least TTP expression level was observed exactly in the samples diagnosed luminal A, which comes into agreement with the previous knowledge.

It was described previously, that expression of ZPF36 can be regulated by methylation [35–37]. It is important regarding to the ability of many of anticancer agents to change the epigenetic landscape [38]. Unfortunately, this type of their action was not studied enough and should be investigated for each tumor suppressor or oncogene individually. The alterations in methylation pattern induced by the therapeutic agents can influence the effectiveness of such therapy and side effects caused by the treatment.

Temozolomide (TMZ) is a DNA alkylating agent of second generation [39]. It is a small lipophilic molecule, the imidazole derivative which hydrolyses to its active intermediate – (3-methyltriazen-1-yl) imidazole-4-carboxamide (MTIC) – at pH >7. One of the described mechanisms of this drug action based on transferring the methyl group to the N7 position of guanine (m^7^G) and N3 position of adenine (m^3^A) [40]. These changes can be repaired in the presence of high levels of MGMT by the base-excision repair mechanism, thus silencing of this enzyme, often by its promoter methylation, increases the effect of TMZ treatment [41]. Although TMZ metabolites transfer methyl groups mainly to purine bases, Barciszewska at al. demonstrated the concentration-dependent induction of Cytosine methylation by this compound [42].

Doxorubicyn (DOX) is an anthracycline antibiotic, widely used in therapy of the number of different cancers. The main characteristics of it as a therapeutic agent is its genotoxisity, caused by intercalation between complementary bases in double-stranded DNA helix and induction of oxidative stress via ROS formation [43]. It is known that oxidative stress induces epigenetic changes, including aberrant hypermethylation of tumor suppressor genes’ (TSGs’) promoters [44].

Our findings show that temozolomid reduces TTPs methylation, which may improve patient prognosis suggesting its onco-suppressing capabilities. On the other hand, doxorobicin promoted TTPs methylation, which may impair its therpeutic effect during breast cancer therapy.

Although further studies with bigger cohort are needed, taking into account high expression level change, we consider that TTP is a potential marker of HER2-enriched breast cancer. Moreover, our findings show that doxorubicine, widely used antitumor antibiotic, increases TTPs methylation, which may impair its therapeutic effect during breast cancer chemotherapy.

## CRediT authorship contribution statement

Experiments on gene expression: Kropyvko S.

Experiments on methylation: Mankovska O.

Statistical analysis and whiting of the manuscript: Hubiernatorova A.

Collection of clinical samples: Syvak L., Verovkina N., Lyalkin S., Lavrynenko K.

Preparation of biological samples: Kropyvko S., Mankovska O., Lavrynenko K., Ivasechko I.

Validation: Stoika R., Rynditch A.

All authors discussed the results and contributed to the final manuscript.

## Acknowledgments

This work was supported by the National Research Foundation of Ukraine Grant (No. 2020.01/0021).

## Declaration of competing interest

The authors of this paper have no conflicts of interest, including specific financial interests, relationships, and/or affiliations relevant to the subject matter or materials included.

## Abbreviations

TTP: Tristetraproline
BC: Breast cancer
RBPs: RNA-binding proteins
AREs: AU-rich elements
qMSP: quantitative methyl specific PCR
TMZ: Temozolomide
Dox: Doxorubicn
ER: estrogen receptor
PR: progesterone receptor
HCC: human hepatocellular carcinoma
TSGs: tumor suppressor genes

## Notes

### Competing Interest Statement

The authors have declared no competing interest.

